# Patient-derived airway organoids from BAL fluid model injury and therapy responses in neonatal bronchopulmonary dysplasia

**DOI:** 10.1101/2025.08.11.669535

**Authors:** Shilpa Sonti, Abiud Cantu, Manuel Cantu Guttierez, Connor Leek, Phinzy Pelton, Erik A. Jensen, Krithika Lingappan

**Affiliations:** Department of Pediatrics, Division of Neonatology, Children’s Hospital of Philadelphia, University of Pennsylvania, PA, USA; Division of Pulmonary and Sleep Medicine, Children’s Hospital of Philadelphia, Perelman School of Medicine, University of Pennsylvania, PA, USA; Department of Pediatrics, Division of Neonatal and Developmental Medicine, Stanford University, Palo Alto, CA, USA

## Abstract

Lung tissue from fetal and neonatal lung samples is tough to obtain, and capturing cells from a living patient with evolving or established disease is very challenging. We hypothesized that airway organoids derived from bronchoalveolar lavage (BAL) samples obtained from intubated preterm infants with bronchopulmonary dysplasia (BPD) will recapitulate the epithelial heterogeneity seen in human airways and can be used to study lung injury and therapeutic response *in vitro*. Here, we demonstrate that BAL sample-derived airway organoids from ventilator-dependent patients with established BPD exhibited cellular heterogeneity consistent with that observed in the human airway. Developed organoids contain basal cell progenitors and a spectrum of differentiated epithelial subtypes, including secretory, ciliated, PNECs, and hillock cells. Hyperoxia exposure and treatment with dexamethasone caused significant cellular transcriptional changes and highlighted biological pathways, both known and novel, with distinct findings based on sex as a biological variable. Findings were validated in an independent dataset from human BPD lung samples. Infant BAL-derived human lung organoids represent a cutting-edge model that bridges a critical gap in BPD research. They combine the advantages of being patient-specific and capturing developmental lung biology, with the experimental flexibility of an *in vitro* system.

## Introduction

Despite significant advances in neonatal care, bronchopulmonary dysplasia (BPD) remains a major cause of morbidity and early mortality among preterm infants. The limited availability of human-specific models that faithfully recapitulate the cellular complexity and developmental context of lung disease in evolving and established BPD contributes to this treatment gap. Lung organoids—three-dimensional structures composed of self-organizing epithelial cells—offer a promising *in vitro* platform to investigate lung development, injury, response to therapy, and repair (*1, 2*).

Lung tissue from fetal and neonatal lung samples is difficult to obtain, and capturing cells from a living patient with evolving or established pulmonary disease poses unique challenges. Conventional organoid systems are typically derived from rare lung specimens (at autopsy or lung biopsy) or induced pluripotent stem cells (iPSCs), each of which presents logistical and technical challenges that limit widespread application. Recent studies have demonstrated that bronchoalveolar lavage (BAL) fluid from adult patients can be used to generate patient-specific lung organoids(3). We hypothesized that airway organoids derived from BAL samples obtained from intubated preterm infants with established BPD will recapitulate the epithelial heterogeneity seen in human airways and can be used to characterize collective cellular function, injury, and response to therapies *in vitro*.

We show that BAL sample-derived airway organoids from patients with established BPD contain basal progenitors and a spectrum of differentiated epithelial subtypes, including secretory, ciliated, PNECs, and hillock cells. We exposed the patient-derived airway organoids to hyperoxia (supraphysiological concentrations of oxygen) and show the resulting transcriptomic responses, including those that varied between male- and female- patient-derived organoids, to highlight the possible disease-modifying role of sex as a biological variable. Finally, we tested the response of the airway organoids to dexamethasone exposure as a model case for using the organoids as a platform to assess therapeutic response and toxicity.

## Results

We used flow cytometry to characterize the cell types present in the airway organoids (**Supplemental Figure 1**). The organoids were enriched in EpCAM+ epithelial cells, and most of them, both in the male and female BAL organoids, were nerve growth factor receptor (NGFR)+ (marker for basal cells). We also sorted NGFR- cells and subjected both to RNA-seq analysis. Differential expression between both cell sub-populations is shown in **Figure 1B**. Expression of genes typically expressed in basal cells, such as *KRT17, KRT5,* and *TP63,* was significantly higher in the NGFR+ cells (**Figure 1C**). The differentially expressed genes in these two cell sub-populations are shown in **Supplemental Table 2.**

**Figure 1:**
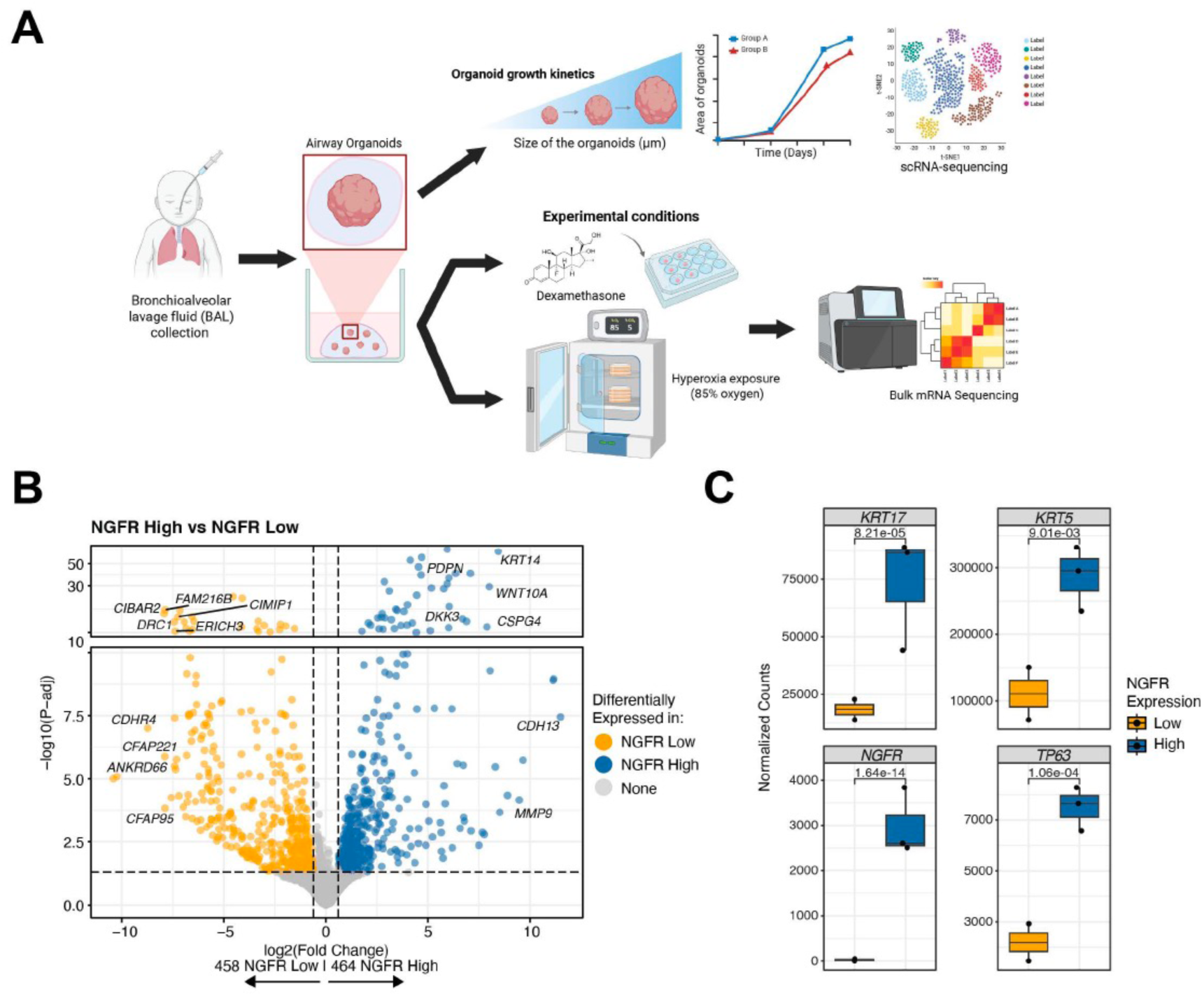
BAL-sample generated Airway Organoids from infants with established BPD are enriched for NGFR+ cells. **(A)** Schematic showing generation of airway organoids from patients with established BPD **(B)** Airway organoids were dissociated and subjected to flow cytometry after 45-70 days of growth. NGFR+ and NFGR- cells were sorted and subjected to flow cytometry. Volcano plot showing differentially expressed genes with fold change on the X-axis and adjusted p-value (FDR) on the Y-axis. **(C)** Expression levels (normalized counts) of *KRT17, KRT5, NGFR, and TP63* in NGFR+ and NGFR- cells show significantly higher expression of basal cell genes in the NGFR+ cells (N = 3/group). Numbers indicate P value for differences between the groups. Created in BioRender. Lingappan, K. (2025) https://BioRender.com/d2jxdgo.

Representative brightfield and IF images of the BAL-derived airway organoids after 14 days in culture are shown in **Figure 2A-C**, respectively. Many cells are positive for TP63 staining and KRT5 (both basal cell markers). We aimed to compare the growth metrics between male and female samples to assess the growth of airway organoids. The organoid-forming efficiency (**Figure 2D**) and the length of the longest axis (after 7 days in culture; **Figure 2D**) were similar between male and female BAL samples. However, the average area (**Figure 2E**) and the average perimeter of the female organoids were significantly higher compared to male organoids at Day 7 and Day 10 of culture.

**Figure 2:**
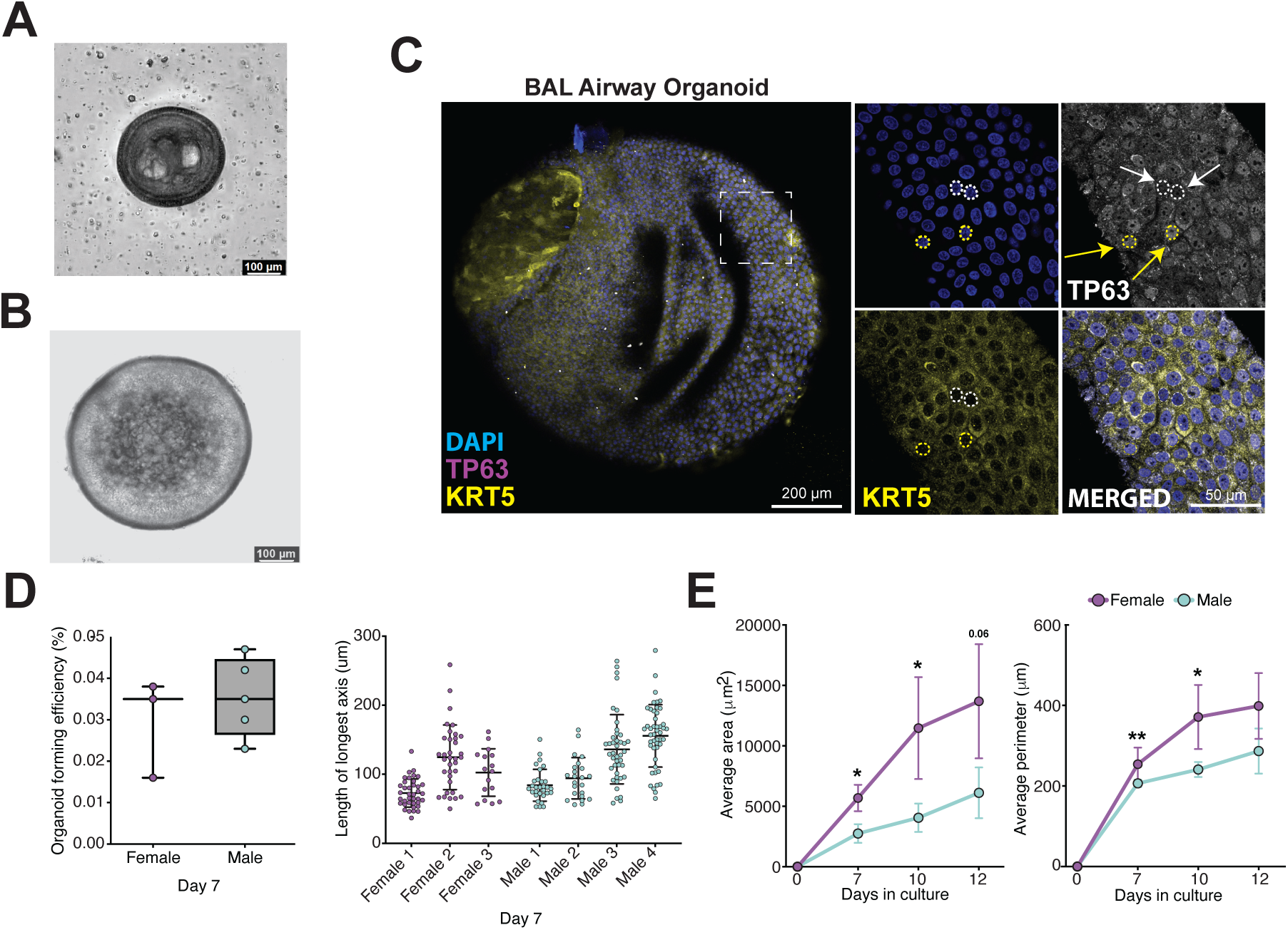
**(A)** Bright field image of male BAL airway organoid taken on day 14 at 4X magnification. **(B)** Bright field image of female BAL airway organoid taken on day 16 at 4X magnification. **(C)** IF image of the airway organoid stained with DAPI, TP63, and KRT5 (basal cell markers). **(D)** Organoid forming efficiency (OFE) of BAL airway organoid samples based on the number of organoids of size >50 μm formed at day 7 following initial seeding of BAL cells. The right panel displays the count and size distribution of organoids formed from each sample on Day 7. **(E)** Organoid growth rate measured in terms of average area and perimeter of the organoids tracked over 12 days. Statistical difference was estimated using Student’s t test; * p<0.05; **p<0.01.

Next, we performed single-cell RNA-Seq on male and female BAL-derived airway organoids to delineate the cellular heterogeneity of the organoids and investigate whether there were any differences based on biological sex. After QC, the data were referenced to previously published human lung single-cell datasets (3–6), and resolved to arrive at biologically meaningful clusters (**Figure 3A-B**), as detailed in the Methods section. Relative abundance of each of the cell types (**Figure 3C**), and the canonical marker genes used to annotate these clusters (**Figure 3D**), are shown. Hillock cells (*KRT13+*) and basal cells made up a significant proportion of the represented cell types. Other cell types included proliferating basal cells (*MKI67+, TOP2A+),* secretory goblet cells (*SCGB1A1, MUC5AB, MUC5AC+),* and PNECs (*ASCL3, FOXI1+)*.

**Figure 3:**
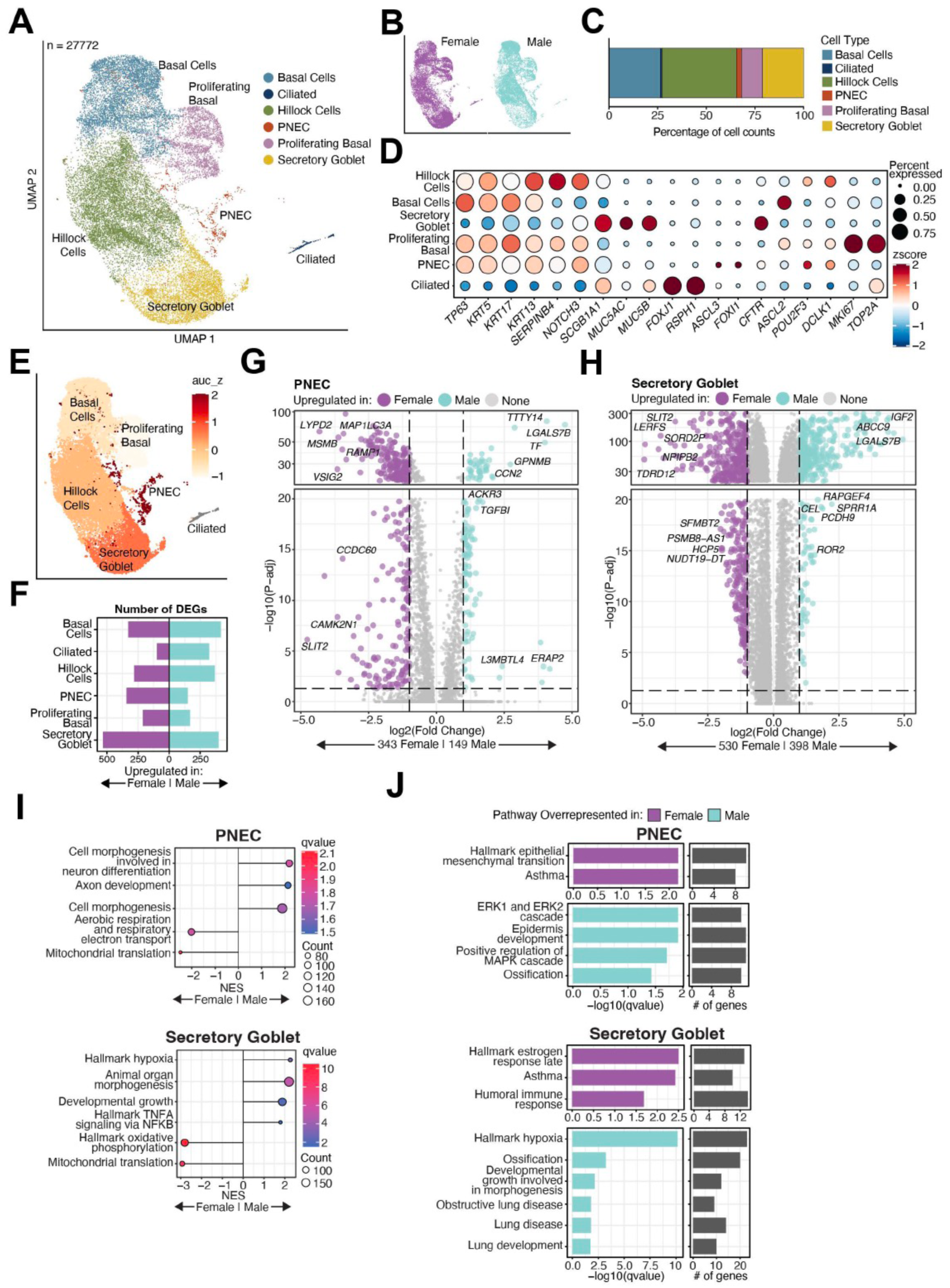
Airway organoids generated from BAL samples from male and female preterm-born infants were subjected to single-cell sequencing after 40-65 days of growth **(A)** UMAP of sequenced cells **(B)** labeled by sex (male or female). Cell clusters were identified based on gene expression. Basal cells, proliferating basal cells, hillock cells, secretory goblet cells, and pulmonary neuroendocrine cells (PNECs) were identified. The total number of cells sequenced is denoted in parentheses. **(C)** The relative % of different epithelial cells comprising the airway organoid. Hillock cells and basal cells were the two main cell sub-populations represented. **(D)** Dot plot showing expression of markers representative of each lung epithelial cell sub-population **(E)** PNECs and Secretory Goblet cells are distinct based on biological sex in BAL-derived airway organoids. Results based on unbiased Augur analysis of the sc-RNASeq datasets. The higher the auc_z score (deeper shade of red), the higher the responsiveness of the cellular transcriptome to biological sex. **(F)** The number of differentially expressed genes between male and female BAL-derived airway organoid cells Volcano plot showing the differentially expressed genes between male and female PNECs **(G)** and secretory goblet cells **(H);** the two cell sub-populations that were most distinct based on biological sex. Pathways analysis in PNECs and secretory goblet cells showing enriched pathways in males and female cells, both by GSEA **(I)** and over-representation (ORA) analyses **(J)**.

Using an unbiased approach to prioritize cell types in the lung most affected by sex as a biological variable, we utilized the Augur bioinformatic pipeline (7). The goal was to identify the cell types most distinct transcriptionally by biological sex. Augur employs a machine-learning framework to rank the cell types found within a single-cell dataset according to the relative magnitude of their response to a biological variable/perturbation. Cell types that are more strongly impacted by the independent variable become more separable within the multidimensional space of molecular measurements than less impacted cell populations. The macro-averaged area under the receiver operating characteristic curve (AUC) for all pairwise comparisons is reported, which provides a single, intuitive measure of cell-type prioritization. An AUC of 0.5 implies that cells cannot be assigned to a specific experimental condition based on their molecular profiles with better-than-random accuracy, and an AUC of 1 reflects perfect classification.

We subjected the sc-RNASeq data sets from the male and female BAL organoids to Augur analysis. We identified that PNECs and secretory goblet cells were the most distinct by biological sex of all the cell sub-populations, with the highest AUC (**Figure 3E**). We then performed differential gene expression analysis of these two cell clusters between the male and female organoids (**Figure 3F-J**). The number of differentially expressed genes between male and female cell sub-populations is shown in **Figure 3F**. DEGs are also visualized in the volcano plot in **Figures 3G and 3H**, which combines the magnitude of change in expression with the significance of the change.

Next, we conducted pathways analysis by GSEA **(Figure 3I)** and over-representation (ORA) analyses to identify biological pathways **(Figure 3J). Supplemental Tables 3 and 4** include the list of DEGs between the male and female cell sub-populations and enriched biological pathways, respectively.

### Hyperoxia exposure leads to a significant alteration of cellular transcription and reflects changes in human BPD lung

Hyperoxia led to strong activation of oxidative stress defenses and cell death pathways, indicating an acute injury response. The number of differentially expressed genes (DEGs), upregulated (410) and downregulated (657) in hyperoxia compared to room air controls, is shown in **Figure 4A**. DEGs are also visualized in the volcano plot in **Figure 4B**, which combines the magnitude of change in expression with the significance of the change.

**Figure 4:**
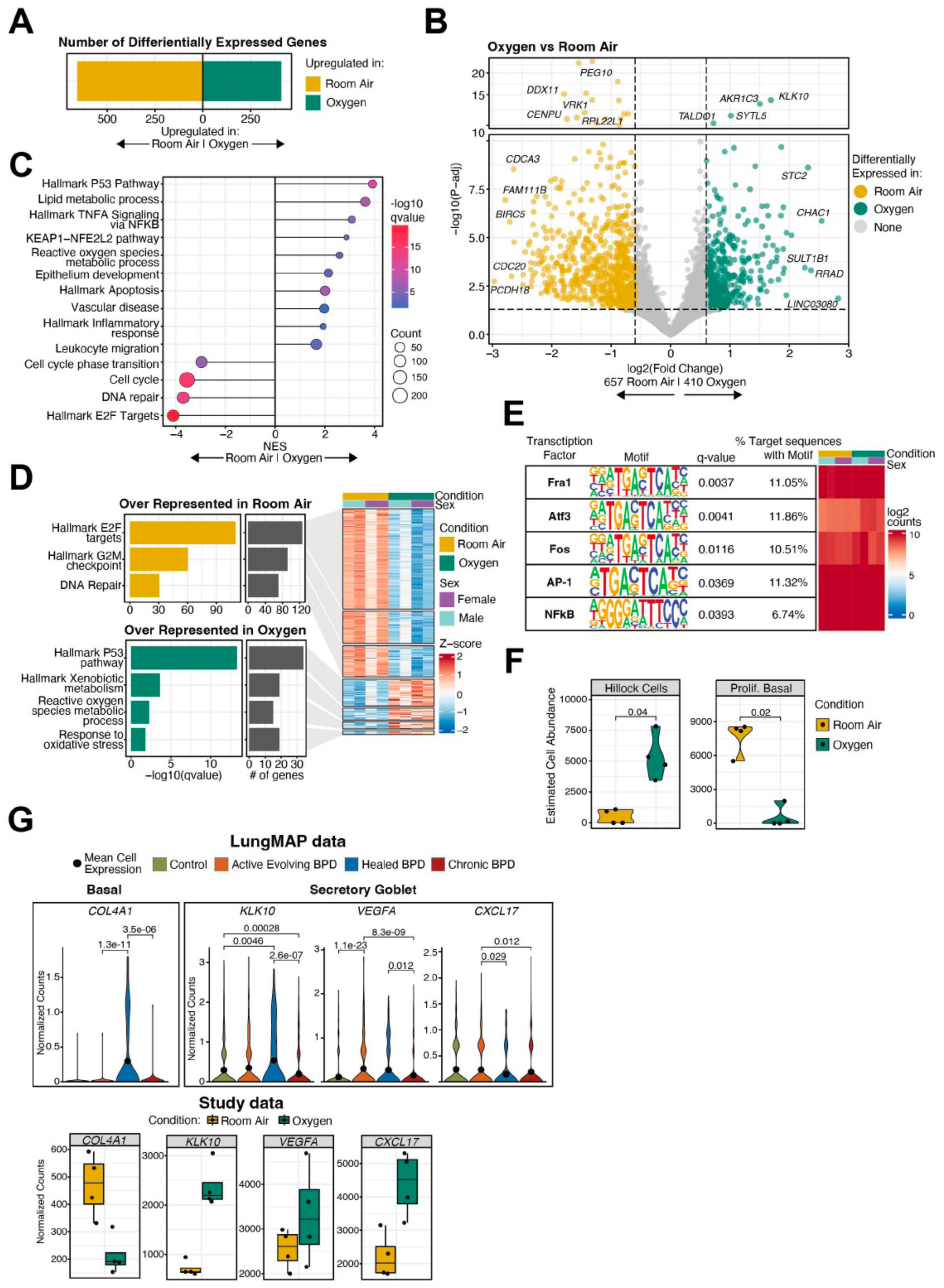
Hyperoxia exposure leads to a significant alteration of cellular transcription and activates apoptosis, inflammation, and pathways related to the metabolism of reactive oxygen species. **(A).** The number of differentially expressed genes (in room air and hyperoxia) in male and female BAL-derived airway organoids (N=4/group; 2/sex/group). **(B)** Volcano plot showing differentially expressed genes with fold change on the X-axis and adjusted p-value (FDR) on the Y-axis. Pathways analysis by GSEA **(C)** and over-representation (ORA) analyses. Component genes from the pathways are represented in the adjoining heatmaps **(D). (E)** HOMER (Hypergeometric Optimization of Motif EnRichment) for motif enrichment analysis to identify overrepresented transcription factor (TF) binding motifs in regulatory regions. We identified potential transcription factors, including *Fra-1, Atf3, Fos, AP-1,* and *NFkB.* (F) Deconvolution analysis suggested that the transcriptional response was being driven by an increase in hillock cells and a decrease in proliferating basal cells in the hyperoxia-exposed organoids. **(G)** Differentially expressed genes in the hyperoxia-exposed BAL organoids were found to be modulated in human BPD (Data from LungMAP). *Genes included COL4A1, KLK10, VEGFA, and CXCL17*.

Next, we conducted pathways analysis by GSEA (**Figure 4C**) and over-representation (ORA) analyses to identify biological pathways. Component genes from the pathways are represented in the adjoining heatmaps (**Figure 4D**). Key antioxidant/detoxification pathways (e.g. NRF2-mediated) are highly enriched, reflecting the response to hyperoxia. Pathways associated with apoptosis and inflammation are also upregulated, consistent with acute lung cell injury under hyperoxic conditions. Cell-cycle gene sets (E2F targets) are markedly downregulated, indicating growth arrest and diminished regenerative proliferation.

Next, to uncover the upstream regulatory mechanisms driving the observed gene expression changes, we used HOMER (Hypergeometric Optimization of Motif EnRichment) for motif enrichment analysis to identify overrepresented transcription factor (TF) binding motifs in regulatory regions. We identified enriched TF motifs in the promoters (or enhancers) of differentially expressed genes (DEGs), helping pinpoint transcription factors likely responsible for the observed gene expression programs. This included transcription factors, such as FRA-1, ATF3, Fos, AP-1, and NF-κ*B (****Figure 4E*).**

To infer the proportion and transcriptional profile of different cell types in our bulk RNA-Seq data from the airway organoids, we utilized single-cell data annotations derived from our organoid samples, as shown in **Figure 2**. Deconvolution allowed us to estimate the relative proportions of different cell types between control (room air) and hyperoxia-exposed airway organoids. Our goal was to identify cell types that may be affected by exposure to hyperoxia. Deconvolution analysis revealed that Hillock cells were estimated to increase, while proliferating basal cells were estimated to decrease upon exposure to hyperoxia **(Figure 4F)**.

We wanted to test if any of the differentially expressed genes in the hyperoxia-exposed BAL-derived airway organoids were captured in the single-cell analysis from human BPD lungs in babies (control, evolving BPD, chronic BPD, and healed BPD). These data were obtained from the LungMAP initiative. Genes differentially modulated in the hyperoxia-exposed BAL organoids were found to be modulated in human BPD, including *COL4A1, KLK10, VEGFA, and CXCL17 (****Figure 4G*).** DEGs and pathways from this study are included in **Supplemental Table 5.**

### Sex as a biological variable modulates the response of the BAL-derived airway organoid to hyperoxia exposure

We subjected male and female BAL-derived airway organoids exposed to room air or hyperoxia to bulk RNA sequencing. The PCA plot **(Supplemental Figure 2A)** showed a clear separation of samples along principal components, validating that both sex and treatment have a strong effect on the cellular transcriptional state. The majority of the differentially expressed genes (DEGs) in male (271 up, 265 down) and female (265 up, 640 down) organoids were unique to each sex, with only 10.3% genes in common **(Figure 5A-B)** between the male and female organoids. DEGs are also shown in the volcano plot in **Figure 5C-D**.

**Figure 5:**
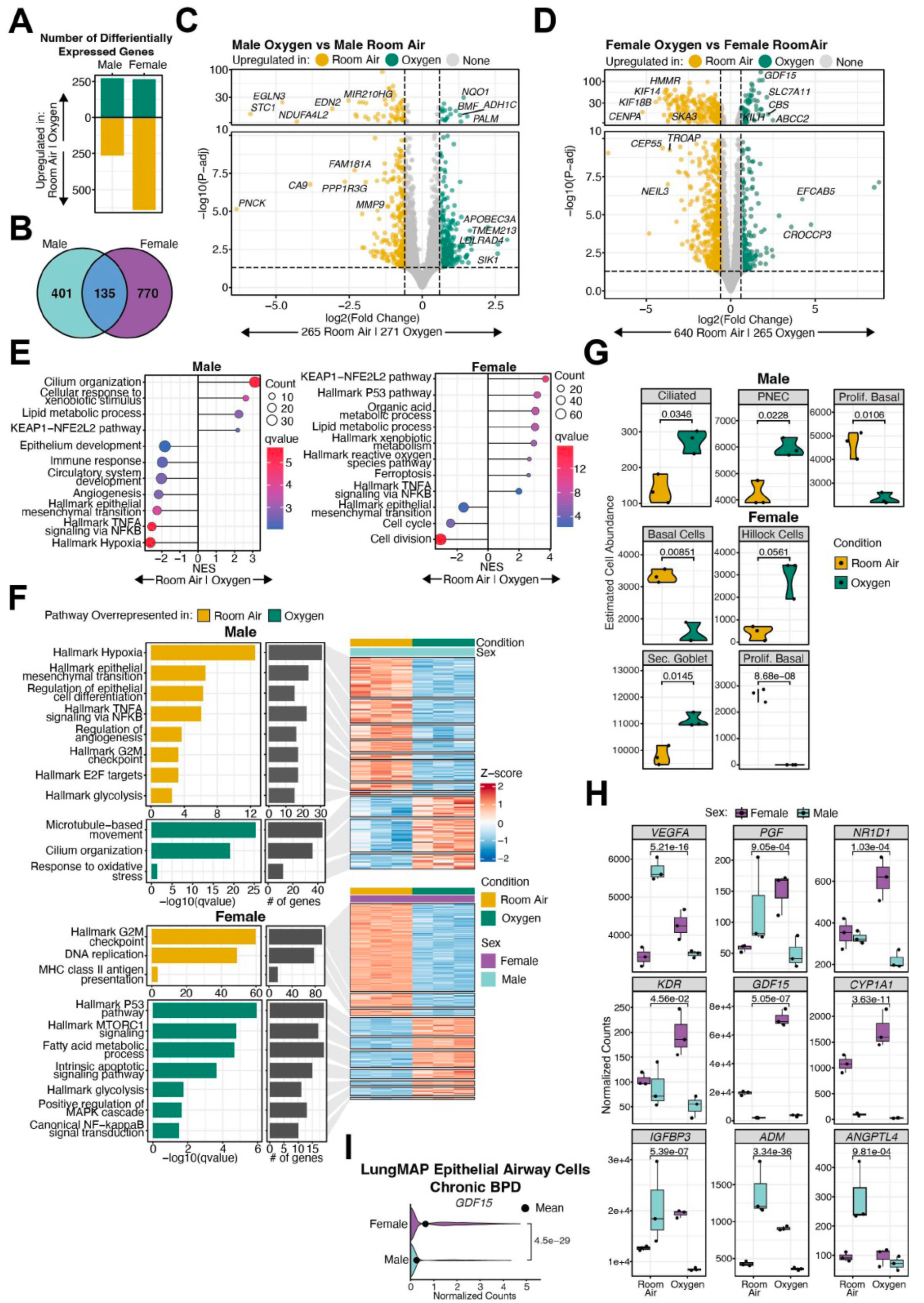
Sex as a biological variable modulates the response of the BAL-derived airway organoid to hyperoxia exposure. **(A)** Number of differentially expressed genes in male and female BAL airway organoids in response to hyperoxia. **(B)** 2-way Venn Diagram showing shared and unique genes between male and female BAL organoids in response to hyperoxia exposure. Volcano plot showing differentially expressed genes in hyperoxia-exposed male **(C)** and female **(D)** airway organoids compared to room air controls. Fold change is represented on the X-axis and adjusted p-value (FDR) on the Y-axis. Pathway analysis by GSEA **(E)** and over-representation (ORA) analyses (**F**). Component genes from the pathways are represented in the adjoining heatmaps in male and female organoids. **(G)** Deconvolution analysis suggested that the transcriptional response was being driven by a decrease in proliferating basal cells in both male and female airway organoids. In male organoids, transcriptional response was driven by an increase in ciliated and PNECs, and an increase in hillock cells and secretory goblet cells in female organoids. **(H)** We performed interaction analysis to identify genes that were differentially modulated in male and female BAL-derived airway organoids exposed to hyperoxia. Several genes with known roles in lung development and repair are shown, including *VEGFA, PGF, NR1D1, KDR, GDF15, CYP1A1, IGFBP3, ADM,* and *ANGPTL4.* Values represent P-values based on comparisons of normalized counts. (I) *GDF15* expression was increased in female airway epithelial cells in chronic BPD in lung samples from preterm neonates. (Data from LungMAP).

To summarize the gene expression changes into coherent biological themes, we performed pathway analysis using both GSEA and ORA analyses. The enriched biological pathways and component genes are shown in **Figures 5E and 5F**. Both sexes exhibited some common responses to hyperoxic stress. Notably, cell cycle arrest was a shared outcome, as both female and male organoids significantly downregulated pathways controlling cell division (G2/M checkpoint, E2F targets, etc.). This indicates that hyperoxia broadly inhibits proliferation regardless of sex. Similarly, both sexes mounted an oxidative stress response, with classic *Nrf2*-driven detoxification pathways (xenobiotic metabolism) being enriched among upregulated genes in both females and males. This suggests a common metabolic adaptation to counteract oxygen-induced oxidative damage. Both sexes also exhibited downregulation of EMT/epithelial remodeling programs, suggesting that hyperoxia tends to preserve the epithelial state (or at least prevent mesenchymal transition) in airway organoids. In summary, suppression of proliferation and enhancement of antioxidant defenses were general features of the hyperoxic response in both female and male cultures.

Despite these commonalities, several pathways were uniquely enriched in one sex but not the other, reflecting sex-specific adaptations. Female organoids uniquely showed up-regulation of inflammatory and immune signaling pathways under hyperoxia – for example, only females had a significant enrichment of TNFα/NF-κB signaling among their up-regulated genes, indicating a female-specific pro-inflammatory response. Females also exhibited enrichment of glycolytic metabolism and specific fatty acid metabolic processes that did not reach significance in males, indicating a greater female tendency to shift metabolism (perhaps to meet energy or biosynthetic demands under stress).

A prominent male-specific response was the induction of ciliogenesis and mucociliary differentiation pathways – numerous cilium assembly/motility gene sets were enriched only in males. This may suggest that male airways respond to hyperoxia by enhancing ciliary function or airway clearance mechanisms, a feature not observed in females. Additionally, angiogenesis-related pathways were significantly suppressed in males (e.g., downregulation of vascular development genes).

We performed deconvolution analysis to infer the proportion and transcriptional profile of different cell types in our bulk RNA-Seq data from the airway organoids. We found that the number of proliferating basal cells was estimated to decrease in both male and female airway organoids. In male organoids, ciliated and PNECs increased, while in female organoids, hillock cells and secretory goblet cells increased (**Figure 5G**).

We performed interaction analysis to identify genes that were differentially modulated in male and female BAL-derived airway organoids exposed to hyperoxia. Interestingly, pro-angiogenic genes, such as *VEGFA, PGF*, and *KDR(VEGFR2)*, were increased in female organoids but decreased in male organoids upon exposure to hyperoxia. *IGFBP3, ANGPTL4*, and *ADM* were decreased in male organoids, while in female organoids, they either remained unchanged or showed a trend towards an increase. *GDF15* and *CYP1A1* showed a female-specific increase in response to hyperoxia **(Figure 5H)**. The LungMAP BPD dataset, comprising preterm infants with BPD and controls, provides high-resolution transcriptomic data from human lungs(6). Data from LungMAP show higher expression of *GDF15* in female airway epithelial cells in chronic BPD, consistent with our observed results in hyperoxia-exposed BAL organoids **(Figure 5I)**. We also analyzed the differences between male and female organoids under room air and hyperoxic conditions. These results are presented in **Supplemental Figure 2.** DEGs and pathways from this study are included in **Supplemental Table 6.**

### Dexamethasone exposure triggers steroid-responsive genes in airway organoids, and sex as a biological variable modulates the cellular transcriptional state

We subjected male and female BAL-derived airway organoids exposed to dexamethasone or vehicle to bulk-RNA sequencing. The PCA plot (**Figure 6A**) showed a clear separation of samples along principal components, validating that both sex and treatment have a strong effect on the cellular transcriptional state. The upset plot **(Figure 6B)** shows the magnitude (number of DEGs) in each comparison and the overlap between groups. The majority of the differentially expressed genes (DEGs) in male (88 up, 132 down) and female (90 up, 112 down) organoids were unique to each sex with 26.3 % genes common between the male and female organoids **(Figure 6C)**. DEGs are also shown in the volcano plot in **Figures 6D-E**. To summarize the gene expression changes into coherent biological themes, we performed pathway analysis using both GSEA **(Figures 6F-G)** and ORA analyses. The enriched biological pathways and the component genes are shown in **Figure 6H**.

**Figure 6:**
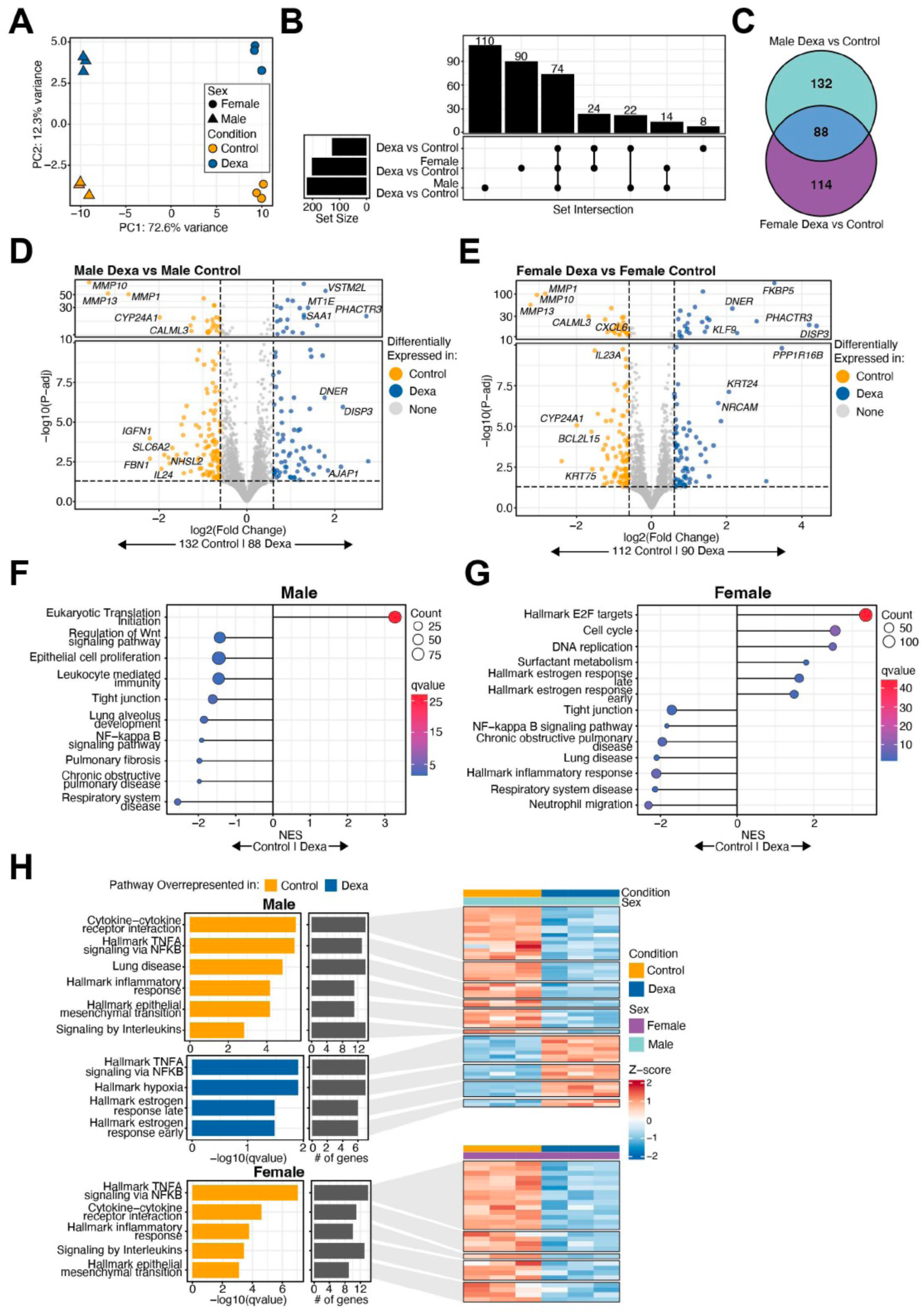
Sex as a biological variable modulates the cellular transcriptional state of the BAL-derived airway organoid after exposure to dexamethasone. **(A)** PCA plot showing separation of the BAL organoids by biological sex (male/female) and treatment (dexamethasone/No Dexamethasone). **(B)** Upset Plot showing the number of differentially expressed genes (DEGs) in male and female BAL airway organoids in response to dexamethasone, showing the lack of overlap DEGs between male and female BAL-derived airway organoids exposed to dexamethasone. **(C)** 2-way Venn diagram showing the common and unique DEGs between male and female BAL-derived airway organoids exposed to dexamethasone. Volcano plot showing differentially expressed genes in dexamethasone-exposed male **(D)** and female **(E)** airway organoids compared to untreated controls. Fold change is represented on the X-axis and adjusted p-value (FDR) on the Y-axis. Pathway analysis by GSEA **(F, G)** and over-representation (ORA) analyses **(H).** Component genes from the pathways are represented in the adjoining heatmaps in male and female organoids, respectively.

Broadly, glucocorticoid (Dexamethasone) exposure suppressed inflammatory/immune and remodeling pathways in both sexes, including the NF-κB signaling pathway and cytokine and chemokine signaling pathways. Female organoids showed strong enrichment of cell cycle (E2F targets, DNA replication), while male organoids showed negative enrichment for epithelial proliferation. Dexamethasone also induced hormone response pathways, such as the Estrogen Response (Early/Late), in both sexes, indicating overlapping gene expression with estrogen-responsive genes. Dexamethasone broadly affected pathways related to tissue remodeling, as both sexes showed significant negative enrichment for the Epithelial– Mesenchymal Transition (EMT) Hallmark gene set, indicating that Dexamethasone suppressed EMT-associated genes. Interestingly, pathways related to tight junctions also appeared with negative NES in both sexes.

Despite the overall overlap, a few pathway enrichments were sex-specific. One striking difference was in lung maturation vs. developmental pathways. Female organoids uniquely showed enrichment of surfactant-associated pathways (e.g., surfactant metabolism positively enriched in GSEA), suggesting that Dexamethasone triggered expression of surfactant proteins (like SFTPB) and related alveolar maturation genes predominantly in females. Correspondingly, female DE gene lists included surfactant protein genes and epithelial sodium channel subunits (*SCNN1A; ENaC*), which are critical for lung fluid clearance at birth – consistent with a Dexamethasone-induced acceleration of functional lung maturation in females. Male organoids, however, did not show significant surfactant pathway enrichment. In fact, GSEA indicated that males had downregulation of some lung developmental gene sets (e.g., lung alveolus development was negatively enriched in males), which was not observed in females. Males, in contrast, showed enrichment of hypoxia-related pathways that females did not.

Dexamethasone-responsive genes were modulated in the male and female BAL-derived airway organoids compared to untreated controls. Examples shown include *FKBP5, ZFP36, NFKBIA, DUSP1, CCL20,* and *PER1* **(Figure 7A)**(8–10). We performed interaction analysis to identify genes that were differentially modulated in male and female BAL-derived airway organoids exposed to dexamethasone. Several genes with known roles in modulating steroid activity and response, as well as lung development, are shown, including *SMC1A, FOXM1, SCNN1A, TSC22D3, TRAF3, THBS1, PDPK1,* and *SFTPB* **(Figure 7B).** Interestingly, LungMAP data showed higher expression of *SCNN1A* in females, and *TRAF3* was higher in male airway epithelial cells in chronic and healed BPD (**Figure 7C**). We also analyzed the differences between male and female organoids after exposure to dexamethasone. These results are discussed and presented in **Supplemental Figure 3.** DEGs and pathways from this study are included in **Supplemental Table 7.**

**Figure 7:**
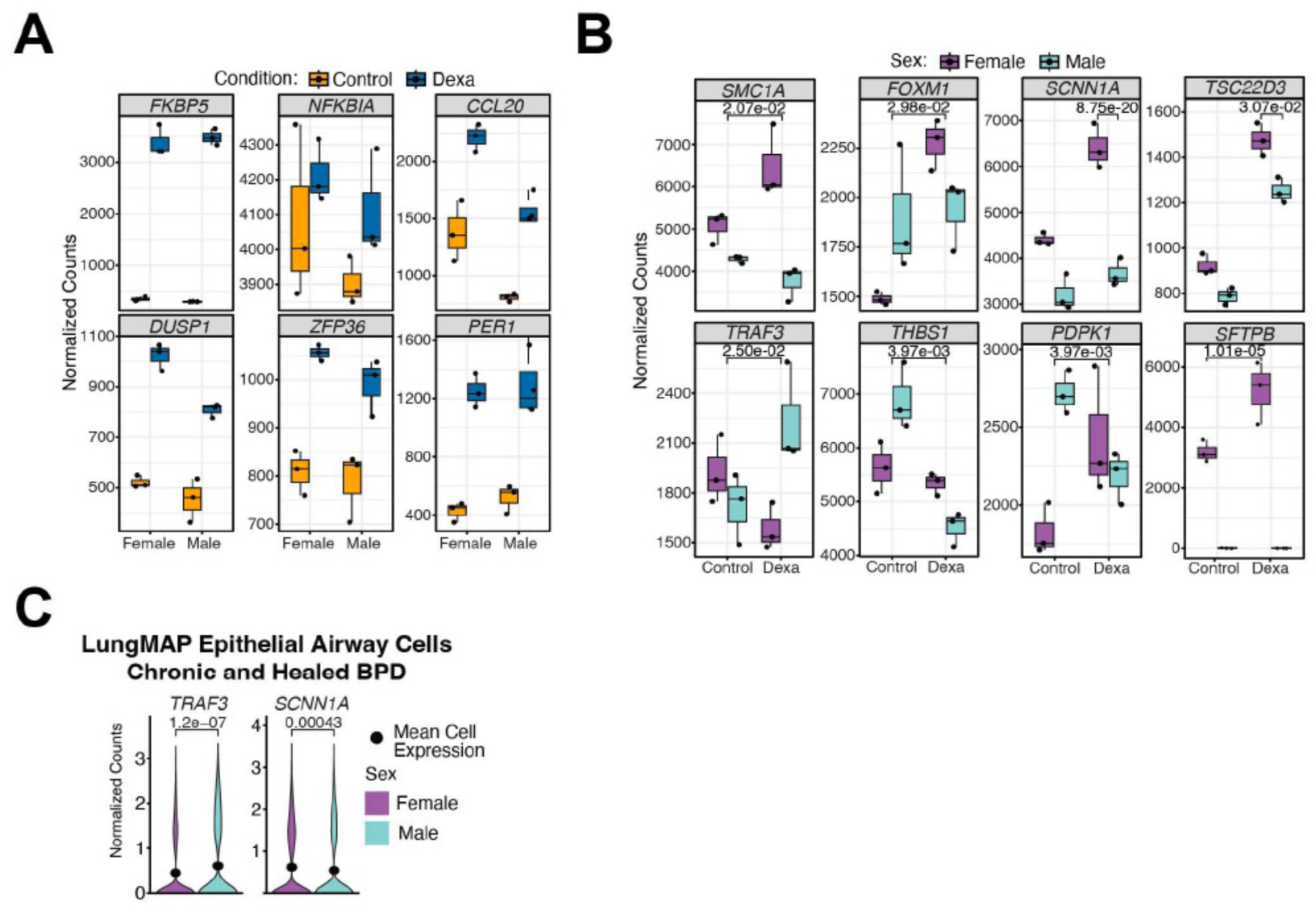
Dexamethasone exposure recapitulates steroid response in patient-derived airway organoids, and sex as a biological variable modulates response to dexamethasone treatment. **(A)** Dexamethasone-responsive genes were modulated in the male and female BAL-derived airway organoids compared to untreated controls. Examples shown include *FKBP5, NFKBIA, CCL20, DUSP1, ZFP36,* and *PER1*. **(B)** We performed interaction analysis to identify genes that were differentially modulated in male and female BAL-derived airway organoids exposed to dexamethasone. Several genes with known roles in lung development and repair are shown, including *SMC1A, FOXM1, SCNN1A, TSC22D3, TRAF3, THBS1,* and *PDPK1.* (C) *TRAF3* expression was increased while SCNN1A expression was increased in female airway epithelial cells in chronic and healed BPD lung samples from preterm neonates. (Data from LungMAP).

## Discussion

Bronchopulmonary dysplasia (BPD) is a chronic lung disease of premature infants characterized by arrested alveolar development and dysregulated repair in the immature lung. A major challenge in studying BPD pathophysiology is the scarcity of accessible human lung tissue from infants with developing or established disease. Traditional animal models (e.g. hyperoxia-exposed neonatal mice, ventilated preterm lambs) replicate certain injury patterns but differ in developmental timing and lung structure, limiting their translational fidelity. Fetal lung explant cultures and precision-cut lung slices preserve 3D architecture and multicellular interactions, allowing tests of injurious stimuli *in vitro.* However, these explant models have short viability and seldom recapitulate the pulmonary vasculature abnormalities that are common in BPD. Given these limitations, new human-derived 3D culture models are being developed to better mimic the complex injury and repair processes of the developing lung.

In particular, advances in stem cell technology and organoid culture offer promising platforms to study BPD’s complex pathogenesis. Lung organoids are 3D spheroid cultures that self-organize from lung epithelial progenitors and recapitulate key structural and functional features of the airway or alveolar epithelium. Traditionally, lung organoids have been generated from pluripotent stem cells or adult tissue biopsies. Recently, researchers have demonstrated that patient-derived cells obtained from bronchoalveolar lavage (BAL) fluid can be utilized to establish both airway and alveolar organoids(3). This breakthrough indicates that even without a surgical biopsy, one can grow patient-specific lung organoids from BAL fluid, capturing both proximal airway and distal alveolar compartments of the lung. We show for the first time that airway organoids derived from BAL samples of patients with established BPD recapitulate the cellular diversity of the human airway, encompassing basal progenitors and multiple differentiated epithelial lineages, including secretory, ciliated, pulmonary neuroendocrine, and hillock cells. The organoids responded to hyperoxia and dexamethasone exposure with marked transcriptional changes, implicating a range of biological pathways—both previously recognized and newly identified—in the pathophysiology of BPD. Importantly, we identified sex as a significant modifier of these transcriptional responses, underscoring the relevance of sex-specific mechanisms in airway injury and therapeutic response. These findings were validated in an independent cohort of human BPD lung specimens, reinforcing the translational relevance of the BAL-derived organoid model.

Our analysis showed that pulmonary neuroendocrine cells (PNECs) and secretory goblet cells were the most distinct based on biological sex. In the lung, sex-dependent neural differences are thought to mediate functional differences related to airway constrictions and mucus clearance (11). A recent study showed that precision-cut lung slices from adult male and female lungs showed distinct neuro-immune outcomes. Another study pointed to a male predominance in the incidence of neuroendocrine cell hyperplasia of infancy (NEHI), a form of childhood interstitial lung disease(12). Not much is known about sex-based differences in the lung secretory goblet cells. Estrogen-mediated increase in mucin secretion from human bronchial epithelial cells has been reported (13).

The genes that were upregulated in the airway organoids were interrogated in the single-cell data from human BPD lungs provided by the LungMAP initiative. We were able to demonstrate the expression of genes such as Kallikrein-Related Peptidase 10 (*KLK10*), Vascular Endothelial Growth Factor A (*VEGFA*), C-X-C Motif Chemokine Ligand 17 (*CXCL17*), and Collagen Type IV Alpha 1 Chain (*COL4A1*) in human BPD in airway epithelial cells. **Table 1** summarizes the known roles of these genes in lung injury.

**Table 1:** Differentially modulated genes in hyperoxia-exposed organoids and human BPD Lungs.

| Table 1: Gene | Role in Lung Injury (Summary) |
| --- | --- |
| **KLK10** (Kallikrein-10) | A secreted serine protease that exerts anti-inflammatory and barrier-protective effects on the pulmonary vasculature, thereby mitigating endothelial dysfunction during lung injury.(14) |
| **VEGFA** (Vascular Endothelial Growth Factor A) | A pivotal regulator of lung vascular and alveolar integrity. In experimental bronchopulmonary dysplasia models, postnatal VEGF supplementation improved survival, enhanced lung angiogenesis, and prevented alveolar damage caused by hyperoxic injury.(15) |
| **CXCL17** (C-X-C Motif Chemokine 17) | A mucosal chemokine highly expressed in the lung, critical for recruiting macrophages and orchestrating immune responses. CXCL17 deficiency leads to fewer lung macrophages and aberrant inflammatory responses to lung insults.(16) |
| **COL4A1** (Collagen, Type IV, Alpha 1) | A key basement-membrane collagen required for alveolar structural integrity. Mutations in *Col4a1* disrupt alveolar septation and the blood-gas barrier, highlighting its essential role in alveolar development and in recovery from lung injury.(17) |

Transcription factors (TFs) likely responsible for the observed transcriptional changes in the airway organoids include FRA-1, ATF3, Fos, AP-1, and NF-κB. The known roles of these genes in lung injury and development are summarized in **Table 2**.

**Table 2:**
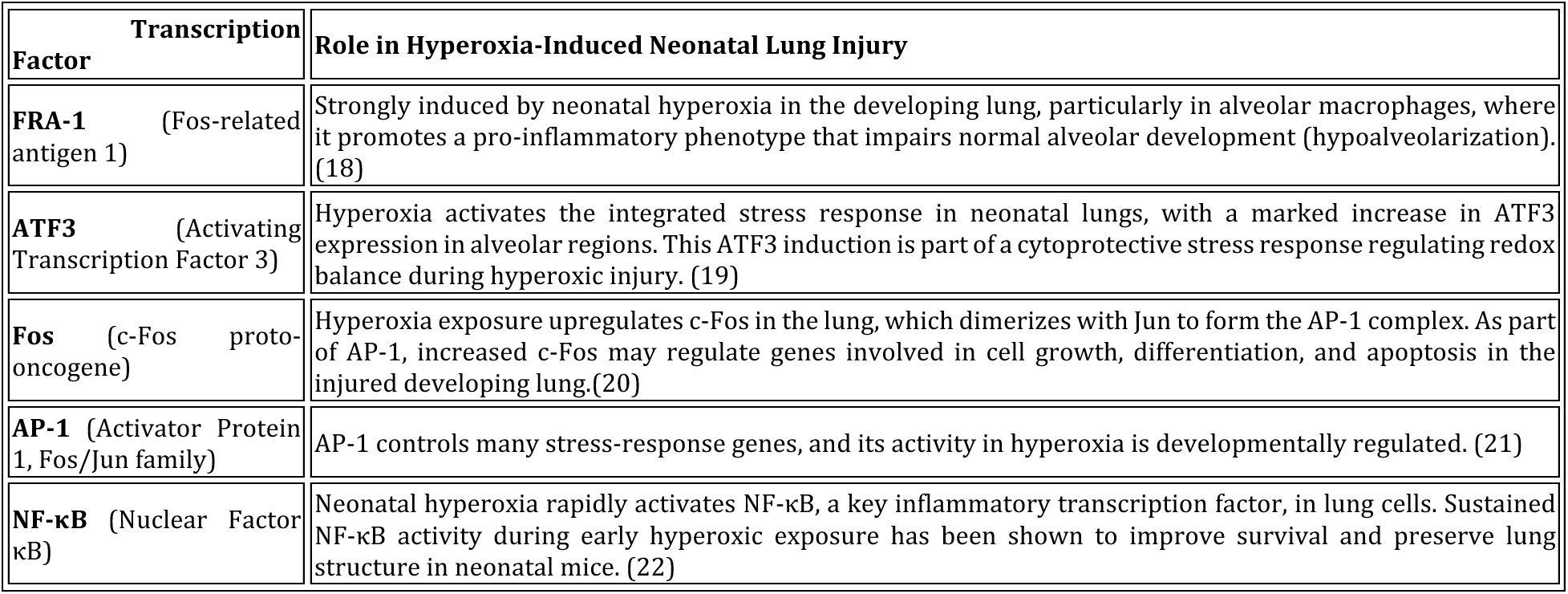
HOMER (Hypergeometric Optimization of Motif EnRichment) to identify transcription factors likely responsible for the observed gene expression programs.

The transcriptomic response in the male and female organoids varied substantially, with the majority of differentially expressed genes being unique to one sex. Both sexes activate protective oxidative stress responses and metabolic adaptations under hyperoxic exposure, while inhibiting cell proliferation and EMT. However, inflammatory and immune signaling emerged as a point of divergence – females tended to enrich for NF-κB pathway activation, whereas males dampened them. Likewise, angiogenesis pathways were only curtailed in males. Males exhibited a unique enhancement of ciliary/airway epithelial remodeling, which was absent in females. These differences suggest that while some core responses to hyperoxia are shared (e.g., antioxidant metabolism, cell-cycle arrest), there are critical sex-specific regulatory differences. Female airway organoids appear to mount a more immune-activated, metabolically active response. These opposing trends suggest that sex can influence the direction of pathway regulation under the same environmental stress, potentially indicating intrinsic differences in regulatory networks or hormone-mediated effects within cells. These findings underscore the significance of sex as a biological variable in airway epithelial stress responses, with implications for differential susceptibility and tissue remodeling in male and female lungs under high oxygen conditions.

Interestingly, deconvolution analysis revealed that female organoids exhibited an increase in hillock cells and secretory goblet cells after hyperoxia. In contrast, male organoids exhibited a rise in ciliated epithelial cells and pulmonary neuroendocrine cells. Hillock cells (23) and ciliated epithelial cells (24) play a crucial role in the regeneration of lung epithelium following injury. Proliferation of neuroendocrine cells (25) after lung injury, and an increase in children with BPD (26), has been reported.

We identified many genes that were modulated differently in the male and female organoids in response to hyperoxia. The list of these genes and their known role in angiogenesis and lung injury is summarized in **Table 3**.

**Table 3:** Differentially expressed genes that were modulated differently in the male and female organoids in response to hyperoxia.

| Table 3: Gene | Direction of Change | Role in Lung Injury (Summary) |
| --- | --- | --- |
| **KDR** Kinase insert domain receptor (VEGFR-2) | ↑ in female | Principal signal-transducing VEGF receptor; enhances endothelial proliferation(27, 28). |
| **GDF15** (Growth Differentiation Factor 15) | ↑ in female | A stress-responsive cytokine that is upregulated in injured lungs; functions as a protective factor in neonatal lung injury. Knockout of *Gdf15* worsens hyperoxia-induced lung damage and impairs alveolar development in neonatal mice.(29) |
| **VEGFA** (Vascular Endothelial Growth Factor A) | ↑ in female | A pivotal regulator of lung vascular and alveolar integrity. In experimental bronchopulmonary dysplasia models, postnatal VEGF supplementation improved survival, enhanced lung angiogenesis, and prevented alveolar damage caused by hyperoxic injury.(15) |
| **IGFBP3** (Insulin-like growth factor binding protein 3) | ↓ in male | Can enhance angiogenesis(30). Recombinant IGFBP-3 lessens lung injury(31). |
| **ANGPTL4** (Angiopoietin-like 4) | ↓ in male | Potent angiogenic factor (32) and has anti-inflammatory activity(33, 34). |
| **ADM** (Adrenomedullin) | ↓ in male | Diminishes lung injury, vasodilatory peptide that up-regulates VEGF (35–37) |
| **CYP1A1**(Cytochrome P450 family 1 subfamily A member 1) | ↑ in female | Protective role against oxidative stress (38) |

Lung organoids offer a physiologically relevant *ex vivo* platform that recapitulates key structural and functional features of the airway epithelium, allowing for the precise evaluation of cellular responses to pharmacological agents. Because organoids retain donor-specific genetic and epigenetic profiles, they offer a powerful model for assessing interindividual and sex-specific variability in drug responsiveness and toxicity. Dexamethasone is widely used in neonatal care to reduce inflammation, promote short-term changes in pulmonary mechanics, and facilitate extubation in preterm infants. However, its potent glucocorticoid activity may disrupt key developmental processes in the immature lung, including alveolarization, vascular growth, and epithelial cell differentiation. Understanding dexamethasone’s effects on the developing lung is essential to balance its short-term respiratory benefits with potential long-term consequences on lung structure, function, and repair capacity (39, 40).

In our experiments, dexamethasone exposure showed anti-inflammatory effects in airway organoids (via NF-κB and cytokine gene downregulation) and showed positive enrichment of surfactant-related pathways. The differences between male and female responses – such as females showing more surfactant upregulation and males showing more hypoxia-related gene activation-suggest that sex-specific factors modulate the response to dexamethasone, which could be important for optimizing clinical steroid use. Overall, these results align with the known actions of glucocorticoids in promoting short-term lung development and controlling inflammation, helping to reinforce our understanding of the therapeutic role of dexamethasone as a prenatal therapy in threatened preterm delivery and in the management of inflammatory lung disease. However, these potential advantageous treatment effects may be counterbalanced by the injurious actions of dexamethasone in the developing lung. While the anti-inflammatory and maturation-promoting effects of dexamethasone are well-established, its suppression of EMT and tight junction gene programs could have unintended consequences on structural lung development, especially if timing, dosage, or duration of exposure is suboptimal (41). We identified many genes that were modulated differently in the male and female organoids in response to dexamethasone. The list of these genes and their known role in response to steroids is summarized in **Table 4**.

**Table 4:** Differentially expressed genes that were modulated differently in the male and female organoids in response to dexamethasone.

| Table 4: Gene | Direction of Change | Response to or Modulation of Steroid Response (Summary) |
| --- | --- | --- |
| **TSC22D3** (GILZ: Glucocorticoid-induced leucine zipper): **X chromosome gene** | F>M | Encodes GILZ, an anti-inflammatory protein. Induced by Dex to mediate glucocorticoid effects – inhibits NF-κB and other pro-inflammatory pathways, thereby suppressing inflammation and promoting resolution.(42) |
| **PDPK1** (3-phosphoinositide-dependent protein kinase-1) | ↑ in female, ↓ in male | Key regulator of the PI3K/AKT signaling pathway, which often intersects with glucocorticoid receptor (GR) signaling.(43, 44) |
| **SCNN1A** (α-ENaC; Epithelial sodium channel α-subunit) | F>M | Encodes the α-subunit of epithelial sodium channel. Dex robustly increases expression in alveolar epithelium.(45) |
| **TRAF3** (Tumor necrosis factor receptor (TNFR)-associated factor 3 tumor necrosis factor receptor (TNFR)-associated factor 3) | ↑ in male, ↓ in female | TRAF3 normally mediates degradation of NF-κB-inducing kinase, thereby restraining non-canonical NF-κB signaling. Inactivation or genetic variants conferring resistance to glucocorticoid therapy.(46–48) |
| **SMC1A** (Structural maintenance of chromosomes 1A) **X chromosome gene** | ↑ in female, ↓ in male | *SMC1A* is a core subunit of the cohesin complex, facilitates glucocorticoid receptor-mediated transcriptional responses by promoting cohesin loading at GR-bound chromatin in a dexamethasone-dependent manner. Patients with cohesin mutations exhibit a reduced response to glucocorticoids(49). |
| **THBS1** (Thrombospondin 1) | ↓ in male | Inhibits angiogenesis in neonatal lungs, upregulates TGF-beta activity(50–52) |
| **FOXM1**(Forkhead box M1) | ↑ in female | Plays an important role in lung development; surfactant homeostasis and lung maturation (53, 54) |

We recognize the limitations of the BAL-derived organoid systems. Organoid cultures from BAL may preferentially expand specific epithelial progenitors while others are under-represented. Another key limitation is that BAL-derived organoids consist primarily of epithelial cells. They lack the mesenchymal, vascular, and immune components that play critical roles in BPD pathogenesis. While organoids form 3D structures, they are usually closed sphere-like cysts of epithelium rather than an intricate alveolar network, thus further engineering is needed to mimic lung architecture and mechanics. Batch-to-batch variability in Matrigel or differentiation protocols can also affect reproducibility.

Despite these limitations, organoids from BAL fluid represent a feasible way to study infant lung cells longitudinally and, in a patient-specific manner. BAL organoids maintain the genetic and epigenetic features of the donor’s cells, providing a disease-specific model of that infant’s lung. This personalized aspect is valuable given the heterogenous nature of BPD development and presentation – organoids can reflect individual variability in vulnerability or response to therapies. Use of BAL samples also allows repeated sampling of preterm infants’ lungs over time with relatively low risk. In 3D culture, BAL-derived organoids can be experimentally exposed to BPD-relevant injuries (e.g., hyperoxia, inflammation, mechanical stretch, infection) to observe their effects on developmental programs.

BAL-derived lung organoids represent a cutting-edge model that bridges a critical gap in BPD research. They combine the advantages of being human, patient-specific, and capturing developmental lung biology, with the experimental flexibility of an in vitro system. These organoids are already shedding light on how the developing human lung responds to injury and could greatly advance our understanding of BPD pathophysiology. Moving forward, refining this model – for example, by incorporating immune cells or blood vessels, and standardizing culture methods – will further enhance its relevance and utility. Together with complementary models like iPSC organoids, lung-on-chip devices, and animal studies, BAL-derived organoids will be a powerful tool in unraveling BPD’s complex biology and in testing novel interventions to protect or regenerate the vulnerable neonatal lung.

## Methods

### Sex as a Biological Variable

Analysis of sex as a biological variable was one of the core objectives of this study. All data were analyzed and reported, taking into account the biological sex of the airway organoid.

### Patient BAL Collection

The study was approved by the Children’s Hospital of Philadelphia Institutional Review Board (IRB protocol # 23-021005). Written informed permission was obtained from the parents or legal guardians of each infant participant before enrollment and sample collection. 500-800µl BAL fluid was collected from 10 infants (Supplemental Table 1) in DMEM + 40% FBS + 1 mM EDTA on ice. 2ml of media was added to reduce viscosity. The sample was filtered using a 40 µm cell strainer and centrifuged at 300 rcf for 10 minutes. Cells were resuspended in 1 mL of media for cell counting (**Figure 1A**). The gestational age, sex, and postmenstrual age at the time of BAL collection are shown in **Supplemental Table 1**. All patients were intubated and receiving mechanical ventilation for severe BPD.

### Establishing organoids from BAL fluid/ lung tissue

Organoids were established using a modified protocol optimized by Liu et al. Briefly, 1 ml of fresh BAL fluid in 1 ml of DMEM + 40% FBS + 1 mM EDTA was centrifuged at 200g for 10 min at 4 °C. Supernatant was collected, and the cell pellet was processed immediately or cryopreserved in Bambanker serum-free freezing media (Bulldog Bio^TM^). For processing, cells were resuspended in 1 ml of DMEM + 40% FBS, and viable cells were counted using Trypan Blue. The cells were then sorted through a 100-µm nylon mesh filter (FisherBrand), and the pellet was collected by centrifuging at 200g for 10 minutes at 4 °C. The cell pellet was typically white; however, if it was visibly red, a red blood cell lysis step was added using RBC Lysis Buffer (Miltenyi Biotech). Cells were resuspended in 1 mL RBC Lysis Buffer and incubated at room temperature for no more than two minutes. 5 mL DMEM + 40% FBS was added to stop the lysis, and the cells were pelleted again to remove the lysis buffer. They were then washed with DMEM + 40% FBS before being seeded for organoid development. Single cells (obtained from BAL) were resuspended in Growth Factor-Reduced, Phenol Red-free, LDEV-free Matrigel (Corning) to a concentration of 100,000 live cells per 50 μL of Matrigel, keeping all materials on ice. Drops of 50 μl each were pipetted onto a prewarmed 12-well plate and allowed to solidify for 25 min at 37°C. Finally, 1 ml of airway medium was added to stimulate the development of airway organoids. Composition of Airway medium – Advanced DMEM/F12, 10 mM HEPES, 1X GlutaMAX, 1X B-27 supplement (Thermo Fisher), 100 μg/ml Primocin (InvivoGen), 1.25 mM N-acetylcysteine, 5 mM nicotinamide (Millipore Sigma), 500 ng/ml R-spondin-1, 25 ng/ml FGF7, 100 ng/ml FGF10, 100 ng/ml Noggin (PeproTech), 500 nM A83-01 (Millipore Sigma), 500 nM SB202190 (Tocris), and 5 μM Y-27632 (ATCC). HRG1-β1 (PeproTech) was added at 50 ng/ml for the first 3-4 days.

### Assessment of Growth metrics

At days 7-10 after the initial plating of BAL cells (passage 0), we assessed the number and size of the organoids (measured based on the longest axis of the organoids >50 μm) yielded for each sample to determine the organoid-forming efficiency (OFE). The OFE was defined as the number of spherical structures (>50 μm) formed at day 7 divided by the total number of cells seeded in each well at day 0. The organoid growth rate was assessed by measuring the average area and perimeter of the organoids over 12 days. Assessment of organoid growth metrics was adapted from previously published protocols (55, 56). Bright field images were taken at 4X and 10X magnification using Leica fluorescence microscope (Leica Biosystems), and image quantification was done using ROI manager option in Fiji (ImageJ).

### Organoid maintenance

Organoids were maintained at 37°C, 5% CO2. Airway or alveolar media was changed every 3-4 days. Organoids were passaged on day 7-10 after initial plating, then every 10-14 days thereafter, depending on the degree of growth. For passaging, the media was replaced with 1 mL of TrypLE Express Enzyme (Thermo Fisher), and the Matrigel dome was disrupted by pipetting once. After incubating for 5 min at 37°C, the suspension was sheared through a pipette five times to dissolve any remaining Matrigel and break up organoids. Cells in TrypLE were diluted 5-fold into DMEM + 10% FBS to stop the dissociation. The pellet was collected by centrifugation at 200g for 10 min, and either resuspended in a desired volume of cold Matrigel and replated or resuspended in Bambanker for cryopreservation. Cells were not counted between passages but split at 1:1-1:4 ratios depending on the organoid confluency and experimental requirement.

### Flow cytometry

For flow cytometry, cells were harvested by incubation in TrypLE at 37°C, with gentle pipetting every 5-10 minutes for up to 40 minutes to achieve a single-cell suspension. Cells were passed through a 40-µm nylon mesh filter (FisherBrand) to remove large clumps, centrifuged at 200g for 10 min, and resuspended in 100 µl FACS Buffer (PBS with 10% FBS, 2 mM EDTA, and 10 µM Y-27632), up to a maximum concentration of 10^6^ cells per 100 µl. Lung cells were stained with mouse IgM anti-human HT2-280 (Terrace Biotech TB-27AHT2-280) at 1:50 dilution for 15 min on ice. Next, goat anti-mouse IgM Dylight 488 (Thermo Fisher SA5-10150) was added at 1:200 dilution for 15 min. Cells were then washed with 1 mL of FACS Buffer and centrifuged at 1000g for 2.5 minutes and then resuspended in 100 μL of FACS Buffer. Resuspended cells were stained 1:50 with the following mouse IgG antibodies for another 15 min on ice: anti-human CD45 APC (BioLegend 304012), anti-human EpCAM PE-Cy7 (BioLegend 324222), and anti-human NGFR APC-Cy7 (BioLegend 345126) and stained 1:50 with viability dye DAPI (Thermofisher 66248) diluted 1:100 in PBS. Cells were rewashed, resuspended in 350 μl FACS Buffer, and analyzed on a 5-laser Cytek Cell Sorter (Cytek).

### Exposure to Hyperoxia/ Dexamethasone

Matrigel drops with confluent growth of organoids were passaged and reseeded in triplicate in a prewarmed 12-well plate per condition. The organoids were allowed to recover for three days. On the day of the experiment, fresh medium was replaced in all organoids. For hyperoxia exposure, one plate was left in the incubator (37 °C, room air) and the other plate was placed in a pre-calibrated ProOX model C21 oxygen chamber (Biospherix) and exposed to hyperoxia (37 °C, 85% oxygen) for 48 hours. For Dexamethasone treatment, Dexamethasone (Millipore Sigma) was added to one set of organoids at a concentration of 250 ng/ml and incubated at 37 °C for 24 hours in room air. Following either exposure, a small portion of the organoid embedded in Matrigel from each condition was collected for immunofluorescence experiments, and the remaining drops were lysed in DNA/RNA shield (Zymo Research) for mRNA extraction, and samples were subject to bulk RNA sequencing.

### Immunofluorescence and immunohistochemistry

The protocol was adapted from a previously published protocol (3). In brief, organoids were released from Matrigel by gentle agitation at 4 °C for 1 hour in cold PBS containing 10 mM EDTA in a 15-ml Falcon tube precoated with 1% BSA. Organoids were left to sink by gravity and were fixed in 10% formalin for 30 minutes at room temperature, followed by three washes with PBS and stored at 4 °C. Using wide orifice pipette tips coated in 1%BSA to prevent sticking, organoids were transferred to a 96 well plate where blocking and permeabilization was done using 0.3% Triton X-100 and 2.5% BSA in PBS for 2 hours at room temperature. The following primary and secondary antibodies were added once at a time and left to incubate overnight at 4C diluted blocking and permeabilization buffer: Rabbit anti-TP63 (1:100, Invitrogen, 703809), Mouse anti-KRT5 (1:50, Invitrogen, MA5-17057), Goat anti-Rabbit Alexa Fluor 488 (1:100, Invitrogen, A-11008), Goat anti-Mouse Alexa Fluor 647 (1:100, Invitrogen, A-21235), Goat anti-Rat Alexa Fluor 647 (1:100, Invitrogen, A-21247). Organoids were then transferred to a custom slide, where a few pieces of tape were used at each end to maintain the organoid structure and mounted using 50% Glycerol. Imaging was done using a Leica DMi8 microscope using a 63x lens.

### Single cell RNA sequencing

Cells were dissociated into single cells by incubation in TrypLE at 37°C for 1-2 hours, with gentle pipetting performed every 5-10 minutes. Cell viability was assessed using Trypan Blue. Cells were passed through a 40-µm nylon mesh filter (FisherBrand) to remove large clumps, then centrifuged at 200g for 10 min. The cells were resuspended in DMEM + 10% FBS to a concentration of 1000-1200 cells/µl. Single-cell libraries were generated at the CHOP Single Cell Technology Core using 10x Genomics Chromium Next GEM Single Cell 3’ v3.1 kits for dual-index, paired-end sequencing. Libraries were sent to Novogene for sequencing at a target of 200,000 reads per cell using an Illumina NovaSeq X Plus. Sequencing data were processed using CellRanger v8.0.1 (10x Genomics) with the GRCh38 human reference genome. The chemistry parameter was set to SC3Pv4 (Single Cell 3’ v4), and the expected cell number was set to 10,000 cells per sample. Initial quality assessment of the raw FASTQ files was performed using FastQC v0.12.1 (57) to evaluate read quality scores, adapter contamination, and overall sequencing metrics. Processed gene expression matrices were imported into R and converted to Seurat objects (v5.1.0) (58) for each sample individually. To address ambient RNA contamination we applied SoupX v1.6.2 (59) with a contamination fraction set to 0.05 (5%).

Cells were flagged as outliers and excluded from analysis if they failed to meet any of the following criteria. For UMI count thresholds, we required a minimum of 1,200 UMIs per cell to exclude empty droplets or cells with insufficient RNA capture. Gene count thresholds were set with a minimum of 500 genes per cell, and a maximum at Q3 + (1.5 × IQR) of the gene count distribution to identify potential doublets or artifacts.

Gene complexity was assessed using the genes per UMI ratio, calculated as log₁₀(genes)/log₁₀(UMIs), which needed to exceed 0.75. This metric identifies cells with low complexity, where few genes are detected relative to total UMI count, indicating potential technical artifacts or dying cells with restricted transcriptional diversity. Additionally, mitochondrial gene expression was calculated using Seurat’s PercentageFeatureSet and restricted to less than 10% of total UMIs. To identify and remove doublets, we employed a dual-filtering approach using both scDblFinder v1.22.0 (60) and DoubletFinder v2.0.4 (61). Both algorithms were run with an expected doublet rate of 7.5%. Only cells identified as doublets by both methods were removed.

Both samples were integrated using Seurat’s reciprocal PCA (RPCA) integration workflow implemented in the ‘IntegrateLayers()’ function. Prior to integration, data were normalized using SCTransform (SCT). Integrated data were subjected to dimensionality reduction using Uniform Manifold Approximation and Projection (UMAP) for visualization. Graph-based clustering was performed using the Louvain algorithm at a resolution of 0.1. Cluster-specific marker genes were identified using Seurat’s ‘FindAllMarkers’ function, which performs differential expression testing between each cluster and all other cells.

Differential gene expression analysis between conditions was performed using Seurat’s ‘FindMarkers’ function with the Wilcoxon rank-sum test. Filtering criteria included: minimum detection rate (min.pct) of 0.1, absolute average log2 fold change ≥1, and adjusted p-value <0.05 using Bonferroni correction for multiple testing. Pathway analysis was done following the same criteria and methods described in the bulk RNA sequencing section.

To annotate cell types in our integrated single-cell RNA-seq dataset, we employed the Universal Cell Embeddings (UCE) framework (62). This approach leverages a foundation model trained on over 36 million cells to generate universal embeddings that capture cell identity across diverse tissues and species. The Seurat-integrated expression matrix and associated metadata were converted to AnnData format for compatibility with the UCE pipeline. We utilized the pre-trained UCE model (33-layer, 8 epochs, 1024 token, 1280 dimension variant) to generate cell embeddings. The parameters used to run the model are: human for species, batch size of 25 cells and a 33-layer transformer. For cell type annotation, we performed supervised label transfer using the Integrated Mega-scale Atlas (IMA) reference dataset provided by the UCE authors. Cell cluster annotations were furthered refined using LungMAP (6).

To identify cell types exhibiting differential responses between experimental conditions, we employed Augur v1.0.3 (63), a computational framework for cell type prioritization in single-cell RNA-seq data (Skinnider et al., 2021). Augur quantifies the separability of cells from different experimental conditions within the transcriptional space of each cell type by training cell type-specific random forest classifiers. For this analysis, we used the integrated and annotated Seurat object containing single-cell RNA-seq data from BAL organoid samples. Cells were stratified by their annotated cell type identities (cluster_name), and we sought to identify cell types showing transcriptional differences between male and female samples (Sex as the experimental label). Augur analysis was performed using the calculate_auc function with raw counts, 50 subsamples, 100 minimum cells per cell type. For each cell type, Augur trained a random forest model to predict the sample sex based on gene expression profiles. The model performance was evaluated using repeated subsampling cross-validation, where the area under the receiver operating characteristic curve (AUC) was calculated for each subsample. Cell types with higher mean AUC values indicate greater transcriptional separability between conditions, suggesting these cell populations are more affected by sex-specific differences.

### Bulk mRNA-Seq analysis

RNA was extracted from organoids exposed to room air – hyperoxia or dexamethasone treatment using Quick RNA miniprep kit (#R1054, Zymo) and shipped to Novogene for bulk RNA sequencing. In brief, RNA integrity was assessed using the Bioanalyzer 2100 system (Agilent Technologies, CA, USA). Messenger RNA was purified from total RNA using poly-T oligo-attached magnetic beads. After fragmentation, the first strand cDNA was synthesized using random hexamer primers. Then the second strand cDNA was synthesized using dUTP, instead of dTTP. The directional library was ready after end repair, A-tailing, adapter ligation, size selection, amplification, and purification. The library was checked with Qubit and real-time PCR for quantification and bioanalyzer for size distribution detection. Libraries were sequenced using Illumina Novaseq X Plus Series. Raw RNA sequencing data integrity was verified through MD5 checksum validation. Quality control, preprocessing, and alignment were performed using the nf-core/rnaseq pipeline (version 3.17.0)(64). Reads were aligned to the mouse reference genome (GRCm39, Ensembl annotation version 111) using STAR(65), and transcript abundance was quantified using Salmon(66). Quality metrics were assessed using MultiQC (67). Transcript abundance estimates were summarized to gene-level counts using the tximport (version 1.34.0) (68) package. The DESeq2 (version 1.46.0) (69) package was used to generate a normalized count matrix, with variance stabilizing transformation applied for visualization and clustering analyses. Low-count genes were filtered prior to differential expression analysis to improve statistical power.

Differential gene expression analysis was performed using DESeq2 (version 1.46.0) (69) to identify significantly differentially expressed genes. Statistical significance thresholds were set at an adjusted p-value < 0.05 and absolute log2 fold change > 0.6. Log fold changes were shrunk using the apeglm method (70) to reduce the impact of genes with low counts and high variability. Functional enrichment analysis was conducted using two complementary approaches: 1) Over-representation analysis (ORA) was performed using clusterProfiler (version 4.14.4)(71) to identify significantly enriched biological pathways and Gene Ontology terms among all and differentially expressed genes. Statistical significance for pathway enrichment was determined using q-value < 0.05. 2) Gene Set Enrichment Analysis (GSEA) was implemented through clusterProfiler using pre-ranked gene lists based on statistical significance and direction of change. This approach considered only genes passing significance thresholds. Pathways with an adjusted p-value < 0.05 were considered significant.

For both pathway analysis approaches, gene sets GO:BP and Kegg were obtained from the clusterProfiler package, Hallmark from Molecular Signatures Database (MSigDB) (version 7.5.1)(72) and Reactome pathways through ReactomePA (version 1.50.0)(73). Transcription factor binding site analysis was performed using HOMER (v4.11.1)(74). Differentially expressed genes were separated into upregulated and downregulated sets, and sequence motifs were identified in promoter regions using findMotifs.pl default parameters. HOMER’s default mouse background sequences were used as controls, and motifs with a p-value < 0.05 were considered significantly enriched. R statistical analyses were performed using R version 4.4.2 within a Bioconductor 3.20 docker container environment.

## Data Availability

The complete bioinformatics workflow and analysis code are available at https://github.com/lingappanlab/2025_Airway_Organoids_From_BAL.

## Author Contributions

SS, AC, MCG, CL, and KL designed the research studies, conducted experiments, acquired data, analyzed data, provided reagents, and wrote the manuscript. EAJ and PP designed the research studies, acquired the patient samples and clinical data, and wrote and reviewed the manuscript.

## Acknowledgements

This work was supported by R01HL168066 to KL and EAJ and R01HL144775 to KL.

